# Efficient primer design for broad-range virus detection: A semi-automated workflow using sequence clustering and varVAMP

**DOI:** 10.64898/2026.09.21.753121

**Authors:** Camille Melissa Johnston, Arianna Mingoli, Anne Mette Hoegh, Louise Lohse, Thomas Bruun Rasmussen

**Affiliations:** Section for Veterinary Virology, Virology & Microbiological Preparedness, Statens Serum Institut, Artillerivej 5, DK-2300 Copenhagen, Denmark; Section for Virus Preparedness, Virology & Microbiological Preparedness, Statens Serum Institut, Artillerivej 5, DK-2300 Copenhagen, Denmark

## Abstract

Broad-range RT-qPCR assays are essential for detecting highly diverse viruses within virus families, but designing suitable primers remains challenging because of extensive sequence diversity and biases in public sequence databases. Here, we present a semi-automated workflow that combines sequence clustering and varVAMP to streamline pan-RT-qPCR primer design. The workflow reduces sequence redundancy, improves the representation of viral diversity, and facilitates the identification of broadly reactive primer candidates. We demonstrate the workflow using the arboviral genera *Orthoflavivirus* and *Alphavirus* as proof-of-concept datasets, showing its applicability to large and genetically diverse viral sequence collections. The workflow provides a practical, scalable, and reproducible framework for developing broad-range primer candidates and may support surveillance, diagnostics, and preparedness for emerging or divergent viruses.

## 1. Introduction

Pan-RT-qPCR assays are essential tools for detecting genetically diverse viruses, enabling broad and sensitive detection across genotypes and lineages, or even at the genus or family level (Escutenaire et al., 2007; Eshoo et al., 2007; Hoffmann et al., 2006; Johnson et al., 2010; Patel et al., 2013; Vijgen et al., 2008). These assays target conserved genomic regions, for example, pan-coronavirus RT-qPCR assays targeting the RNA-dependent RNA polymerase (RdRp) gene. Targeting conserved regions ensures robustness despite rapid viral evolution and is particularly important for identifying emerging or re-emerging viruses for which strain-specific assays may fail. Pan-assays are therefore particularly valuable as screening tools, allowing broad detection before more targeted approaches are applied.

Designing broadly reacting primers can be challenging for viruses with high genomic variability, as primers must target conserved regions with minimal sequence variation, while avoiding insertions or deletions. Degenerate nucleotides can increase primer inclusivity, but their use must be balanced against potential losses in specificity and amplification efficiency. varVAMP enables the automated design of degenerate primers for (RT)-qPCR using only a multiple sequence alignment (MSA) as input (Fuchs et al., 2025). However, varVAMP assumes that the MSA accurately reflects viral diversity, which is often not the case due to biases in public repositories. These include the overrepresentation of human-pathogenic strains, the underrepresentation or absence of rare or recently identified variants, and uneven geographic representation of circulating lineages (Chen et al., 2022; Cobbin et al., 2021; Kieft and Anantharaman, 2022; Ritsch et al., 2023). Careful sequence curation and data reduction are therefore essential. Furthermore, the ability of varVAMP to identify suitable primer sets depends strongly on the user-defined identity threshold and maximum number of allowed ambiguous bases. Consequently, systematic optimization of these parameters is recommended (Fuchs et al., 2025).

Emerging viruses, including zoonotic and vector-borne pathogens, continue to pose significant threats to public health, global economies, and wildlife conservation, leading to recurrent outbreaks of international concern (Bhatt et al., 2013; Chancey et al., 2015; Gubler, 2002; Mayer et al., 2017; Qin et al., 2024; Ribeiro dos Santos et al., 2025). Among these, members of the genera *Orthoflavivirus* and *Alphavirus* are arthropod-borne, single-stranded RNA viruses that have caused substantial disease burdens and public health challenges over the past century due to their broad geographic distribution, vector-mediated transmission, and continued emergence into new geographic regions (Føh et al., 2026; Gelskov et al., 2026; Weaver, 2013; Weaver and Reisen, 2010).

The genus *Orthoflavivirus*, within the family *Flaviviridae*, comprises more than 70 small, enveloped, arthropod-borne viruses, several of which are major human pathogens. Notable mosquito-borne orthoflaviviruses include the hemorrhagic fever viruses dengue virus (*O. dengue*; DENV) and yellow fever virus (*O. flavi*; YFV), and the neurotropic viruses Zika virus (*O. zikaense*; ZIKV), West Nile virus (*O. nilense*; WNV), Usutu virus (*O. usutuense*; USUV), Japanese encephalitis virus (*O. japonicum*; JEV), and Saint Louis encephalitis virus (*O. louisense*; SLEV). Tick-borne members include tick-borne encephalitis virus (*O. encephalitidis*; TBEV), Kyasanur forest disease virus (*O. kyasanurense*; KFDV), and Powassan virus (*O. powassanense*; POWV) (Becker et al., 2012; Rathore and St. John, 2020; Simmonds et al., 2025; Zhao et al., 2021).

The genus *Alphavirus*, the sole genus within the family *Togaviridae,* currently comprises 32 recognized species, the majority of which are arthropod-borne (Forrester et al., 2023; Simmonds et al., 2024; Walker et al., 2019). Many alphaviruses are important pathogens of humans and domestic animals. The New World alphaviruses, including Venezuelan equine encephalitis virus (*A. Venezuelan*; VEEV), Eastern equine encephalitis virus (*A. eastern*; EEEV), and Western equine encephalitis virus (*A. western*; WEEV), primarily cause encephalitic disease, whereas the Old World alphaviruses, such as Ross River virus (*A. rossriver*; RRV), Barmah Forest virus (*A. barmah*; BFV), O’nyong-nyong virus (*A. onyong*; ONNV), Chikungunya virus (*A. chikungunya*; CHIKV), and Sindbis virus (*A. sindbis*; SINV), are predominantly associated with febrile illness and arthralgia syndromes (Tsai et al., 2002).

Here, we present a workflow that integrates varVAMP with upstream database construction and sequence clustering, followed by downstream post-processing (**Figure 1**). We applied this workflow to two different viral genera: *Orthoflavivirus* and *Alphavirus*, to generate pan-RT-qPCR assays. Our results demonstrate the utility of this workflow for designing broadly reactive primers for virus detection.

**Figure 1.**
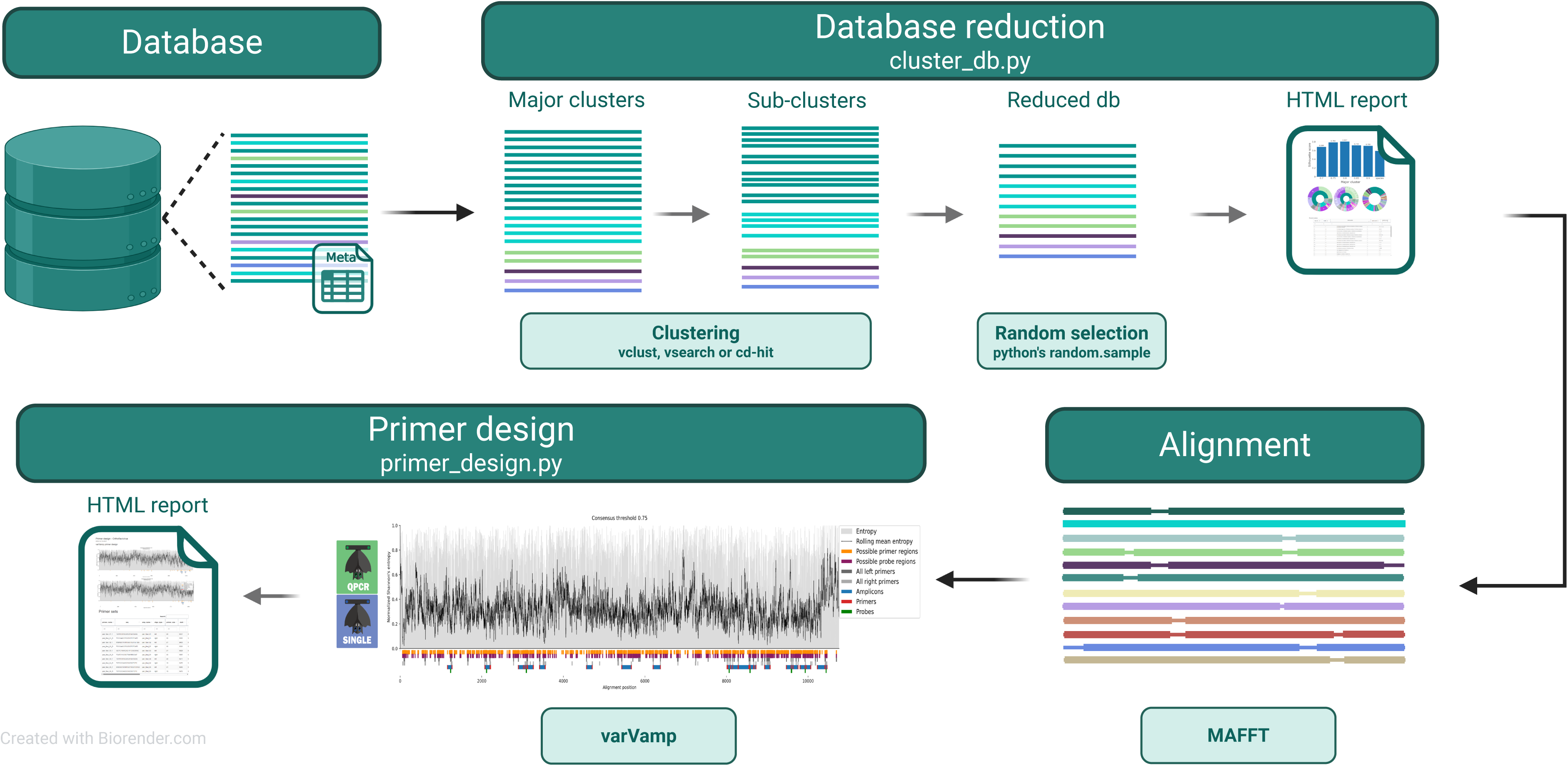
Overview of the semi-automated workflow for pan-primer design. An input sequence database and associated metadata are subjected to hierarchical database reduction, in which sequences are first divided into major clusters and subsequently into subclusters using Vclust, VSEARCH, or CD-HIT. Representative sequences are selected from the resulting subclusters to generate a reduced database, and the clustering and database-reduction results are summarized in an interactive HTML report. The reduced database is aligned using MAFFT and subsequently used as input for primer design with varVAMP. Multiple combinations of primer design parameters are evaluated, and the resulting candidate primer sets are collated and visualized in a separate HTML report for downstream evaluation. Created in BioRender. Johnston, C. (2026). https://BioRender.com/j924ywt

## 2. Methods

### 2.1. Database preparation

All sequences longer than 9,000 nt belonging to the genera *Orthoflavivirus* and *Alphavirus* were downloaded from NCBI (accessed October 2, 2024 and April 20, 2025, respectively) as GenBank files with associated metadata. Entries containing the terms “vector”, “synthetic”, “replicon” or “chimeric” in their titles were excluded. Duplicate sequences were removed, and only sequences with ≤1% ambiguous nucleotides were retained. To restrict the datasets to near-complete or complete genomes, sequences were further filtered by genome length, retaining those between 9,000 and 12,000 nt for *Orthoflaviviris* and between 9,000 and 13,000 nt for *Alphavirus*. The resulting curated datasets were used as input for the clustering workflow.

### 2.2. Clustering and database reduction

The clustering workflow operates through a two-stage hierarchical process, in which viral genomes are first grouped into major clusters and subsequently divided into subclusters. In the first stage, sequences are grouped into major clusters based on user-defined sequence identity thresholds (major cluster IDs). In the second stage, each major cluster is further partitioned into corresponding subcluster identity thresholds (subcluster IDs). The workflow requires a FASTA file containing all the sequences and TSV metadata file specifying sequence accession numbers and species-level taxonomic assignments. Multiple combinations of major cluster and subcluster identity threshold can be specified, generating a corresponding set of clustered and reduced databases.

The workflow supports several clustering back-ends that follow the same general clustering scheme but differ in their implementation and computational characteristics. CD-HIT (Li and Godzik, 2006) uses a fast, greedy incremental clustering algorithm, in which sequences are sorted by length and each assigned to the first existing cluster whose representative (centroid) meets the specified sequence identity threshold. However, *cd-hit-est* is limited to identity thresholds of ≥80%. VSEARCH (Rognes et al., 2016) employs a centroid-based, greedy clustering strategy based on pairwise sequence alignments, allowing more accurate clustering at lower identity thresholds and reducing sensitivity to input order. Vclust (Zielezinski et al., 2025) was later integrated to enable clustering based on average nucleotide identity (ANI). It constructs an all-vs-all ANI matrix, which can be reused to generate cluster partitions across multiple similarity thresholds. Vclust provides six clustering algorithms: single-linkage, complete-linkage, UCLUST (Edgar, 2010), CD-HIT (greedy incremental), greedy set cover (adopted from MMseqs2 (Steinegger and Söding, 2017)) and Leiden (Traag et al., 2019) algorithm. UCLUST, greedy set-cover and CD-HIT are inherently threshold-based algorithms, single and complete linkage algorithms construct dendograms (tree-like structures that represent hierarchical relationships among sequences) that can be pruned by customizable distance thresholds (Zielezinski et al., 2025).

Following subclustering, approximately *n* target sequences (user-defined) are randomly selected from each major cluster using Python’s *random.sample*, while ensuring that at least one representative sequence from each subcluster is retained. Consequently, more than *n* sequences can be selected when the number of subclusters within a major cluster exceeds *n*., while all sequences are retained for clusters containing fewer than *n* sequences.

The outputs are automatically collated into an HTML report summarizing the clustering outcomes. The report includes sunburst plots structured according to species-level metadata, illustrating hierarchical relationships among species, major clusters, and subclusters, as well as the composition of the resulting reduced databases. Each visualization is accompanied by an interactive DataTable containing the corresponding data for inspection. The selected reduced database is subsequently extracted for downstream analysis.

During initial workflow development, selection of the reduced database for downstream analyses was performed empirically based on qualitative assessment of the HTML reports. This approach was subsequently supplemented with a quantitative scoring system to enable objective evaluation of clustering and database-reduction outcomes.

### 2.3. Scoring system for cluster evaluation

A two-level scoring system was implemented to provide an objective framework for evaluating clustering performance. At the major clustering level, silhouette scores (Rousseeuw, 1987), ranging from -1 to 1, were calculated using distance matrices derived from either Mash distances (Ondov et al., 2016) or converted total ANI (tANI) values generated by Vclust. The silhouette scores provide a numerical measure of overall clustering quality by quantifying both intra-cluster cohesion and inter-cluster separation, with values approaching 1 indicating well-separated clusters, values near 0 indicating overlapping clusters, and negative values indicating that sequences may have been assigned to inappropriate clusters. providing a numerical measure of overall clustering quality.

At the subclustering level, an overclustering penalty was applied to penalize excessive fragmentation of major clusters. The penalty was derived from the user-defined *n* target sequences per major cluster in the reduced database. For a given major cluster, if the number of resulting subclusters (*C*) was less than or equal to the target number of sequences (*n*), the overclustering penalty (*D*) was set to 0. Otherwise it was calculated as:

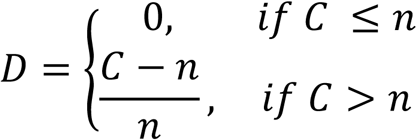

To summarize subclustering performance, two aggregate measures were calculated: an unweighted mean penalty score and a weighted mean penalty score. The unweighted penalty was calculated as the arithmetic mean of the individual penalties across all major clusters, giving each major cluster equal contribution irrespective of its size. The weighted penalty accounts for cluster size and was computed as:

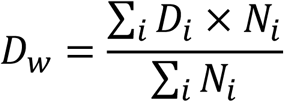

Where *Di* is the overclustering penalty for major cluster *i* and *N_i_* is the number of sequences within that major cluster. Thus, major larger clusters contribute proportionally more to the weighted penalty than smaller clusters.

As the overclustering penalty is directly influenced by the user-defined *n* target sequences per major cluster, careful selection of this parameter is important. The value of *n* should be considered in relation to the size and composition of the major clusters and the selected subclustering threshold, as these factors together determine sequence representation in the reduced database. For example, a higher subclustering identity threshold may divide major clusters into a larger number of subclusters, potentially resulting in more representatives than the specified *n*. In addition, when biologically important groups are represented by substantially fewer sequences than highly sampled groups, selecting an *n* closer to the size of these underrepresented groups may provide a more balanced representation of genetic diversity in the reduced database.

### 2.4. Computational benchmarking

Cluster workflow performance was benchmarked on a dedicated Linux server equipped with an Intel Xeon Platinum 8380 CPU (2.30GHz), running Rocky Linux 9, and managed through the SLURM workload manager. Jobs were executed via *srun* with an allocation of 10 CPUs and 128 GB RAM. All workflows were run using identical thread allocations to ensure comparability. Runtime and memory usage were recorded using GNU */usr/bin/time -v* utility, which reports wall-clock time and peak resident memory. Each benchmark was performed three times and mean values were used for comparison. The cluster workflows were run using major cluster IDs 0.70-0.90 in 0.05 increments, with subcluster IDs 0.96-0.99 in 0.01 increments.

### 2.5. Primer design workflow

The primer design workflow takes a multiple sequence alignment of the reduced database as input, which can be generated using any compatible alignment tool. Here, MAFFT (Katoh and Standley, 2013) with default parameters was used to generate the input alignments. The workflow allows users to specify multiple consensus thresholds and maximum numbers of allowed ambiguous bases (max_ambig). Primer design can be performed in two modes: qPCR and single. In qPCR mode, varVAMP attempts to design primer-probe sets, whereas single mode generates primer pairs without probes. If qPCR mode is selected but varVAMP fails to generate suitable primer-probe sets, the workflow automatically switches to single mode. Outputs for each combination of consensus threshold and max_ambig setting are summarized in an HTML report which visualizes primer coverage and genomic locations and includes regenerated varVAMP overview plots of the predicted amplicons.

### 2.6. Prediction of on-target and off-target hits

Inspired by the amplicon off-target prediction implemented in varVAMP, potential on-target and off-target binding of primer candidates can be evaluated using BLAST (Camacho et al., 2009) against custom databases. Primer candidates are first expanded into all possible ambiguous sequence variants, which are subsequently queried against the respective databases using BLASTn. The resulting alignments are filtered according to criteria specific to on-target and off-target prediction. For both predictions, hits are retained only when the three terminal nucleotides at the 3′-end match perfectly, ensuring compatibility with priming requirements. For on-target prediction, additional filters are applied: (i) the alignment covers at least 75% of the primer length, (ii) the number of mismatches does not exceed the user-defined mismatch tolerance, (iii) the predicted amplicon length falls within a dynamically calculated range of the expected product size, with the allowable deviation scaled according to amplicon length, and (iv) directionality is enforced (e.g., forward primers must align in the forward orientation). For off-target prediction, less stringent filtering criteria are applied: (i) the alignment covers at least 50% of the primer length, (ii) the number of mismatches does not exceed one per five nucleotides, and (iii) the predicted amplicon length (X) should fall within an acceptable range (0.5 × amplicon length ≤ X ≤ max(1000 bp, 1.5 × amplicon length)).

### 2.7. *Orthoflavivirus* RT-qPCRs

A representative panel of *Orthoflavivirus* RNA samples (**Table S1**) was tested using an established pan-flavivirus RT-qPCR assay (Patel et al., 2013) with a modified protocol, to obtain reference Ct-values. Reactions were performed using the Qiagen OneStep RT-PCR kit (QIAGEN, Hilden, Germany) with Resolight Dye (Roche, Basel, Switzerland) in a final reaction volume of 12.5 µL containing 0.5 µM of each primer. Cycling conditions consisted of 50°C for 30 min, followed by 95°C for 15 min, then 40 cycles of 95°C for 15 s, 56°C for 30 s and 72°C for 30 s. Amplication was followed by melting curve analysis.

Primers targeting *Orthoflavivirus* and generated using the workflow were subsequently evaluated by RT-qPCR using the same viral RNA panel. Reactions were performed using the same reaction composition and cycling conditions as described above.

### 2.8. *Alphavirus* RT-qPCRs

A representative panel of *Alphavirus* RNA samples (**Table S2**) was tested using an established pan-alphavirus RT-qPCR assay (Eshoo et al., 2007), using a modified protocol to obtain reference Ct values. Reactions were performed using the Qiagen OneStep RT-PCR kit with Resolight Dye in a final reaction volume of 12.5 µL containing 0.25 µM of each primer. Cycling conditions consisted of 50°C for 30 min, followed by 95°C for 15 min; 8 cycles of 95°C for 30 s, 48°C for 30 s, and 72°C for 30 s, with the annealing temperature increasing by 0.9°C per cycle; followed by 37 cycles of 95°C for 15 s, 56°C for 20 s, and 72°C for 20 s. A final extension was performed at 72°C for 2 min, followed by melting curve analysis.

Primers targeting *Alphavirus* and generated using the workflow were evaluated by RT-qPCR. Candidate primer sets from the initial workflow were first screened using 10,000-fold dilutions of VEEV and CHIKV RNA samples. Primer pairs that successfully amplified both targets were subsequently evaluated against the full, undiluted *Alphavirus* RNA panel. Following reanalysis of the sequence database using Vclust-based clustering, a second set of candidate primer pairs was designed and evaluated. These primers pairs were again screened using 10,000-fold dilutions of VEEV and CHIKV RNA samples, after which selected primer pairs were evaluated against a 100-fold dilution of the *Alphavirus* RNA panel. All workflow-generated primer pairs were evaluated using the Qiagen OneStep RT-PCR kit with Resolight Dye in a final volume of 12.5 µl, containing 0.5 µM of each primer. Cycling conditions consisted of 50°C for 30 min, followed by 95°C for 15 min, then 45 cycles of 95°C for 15 s, 58°C for 30 s and 72°C for 30 s. Amplification was followed by melting curve analysis.

## 3. Results

### 3.1. Database composition and initial reduction of the viral databases

#### 3.1.1. Dataset preparation and initial clustering workflow

Genomes belonging to *Orthoflavivirus* and *Alphavirus* were downloaded from NCBI as described above. For *Orthoflavivirus,* 17,644 sequences were retained after filtering, of which more than 70% belonged to DENV (**Table S3**). For *Alphavirus*, 4270 sequences were retained, of which more than 56% belonged to CHIKV (**Table S4**). Both databases were subjected to the clustering workflow using major cluster ID thresholds of 0.70, 0.75, 0.80, 0.85, and 0.90. VSEARCH was used at ID thresholds below 0.80 and CD-HIT at thresholds of 0.80 and above. Subclustering was performed using ID thresholds of 0.97, 0.98, 0.99, and 0.995 with CD-HIT, with a target of *n*=100 sequences per major cluster.

#### 3.1.2. Clustering and reduction of the *Orthoflavivirus database*

At a major cluster ID of 0.70, species were not clearly delineated, indicating insufficient sequence separation at this threshold. At 0.75, most *Orthoflavivirus* species formed distinct clusters, with the exception of several multi-species clusters: cluster 0 contained TBEV, *O. loupingi*, and *O. omskense*; cluster 14 contained *O. bagazaense*, *O. israelense*, and *O. ntayaense*; cluster 15 contained *O. koutangoense* and WNV; and cluster 44 contained *O. sepikense* and *O. wesselbronense* (**Table S5**). Additionally, WNV was divided into three clusters, while DENV formed four (**Table S6**).

At a major-cluster identity threshold of 0.80, species-level delineation improved further. *O. omskense* no longer grouped with TBEV and *O. loupingi*, and *O. ntayaense* no longer clustered with *O. bagazaense* or *O. israelense*. *O. koutangoense* remained associated with one sequence of WNV. WNV was divided into seven clusters, although most sequences were contained within two major clusters. Similarly, *O. sepikense* and *O. wesselbronense* also formed distinct clusters, while DENV was divided into eight clusters, with the majority of sequences contained within four dominant clusters.

Based on the observed species-level delineation, a major-cluster identity threshold of 0.80 was selected for subsequent analyses. Using a subcluster identity threshold of 0.97, the database was reduced to 1,971 sequences, whereas at 0.98 resulted in 2,009 sequences. The difference between these reductions was primarily attributable to an increased number of DENV subclusters at the higher identity threshold. As this difference was considered negligible, the reduced database generated using a major cluster identity of 0.80 and a subcluster identity of 0.98 was selected for downstream alignment (**Table S7**). In this reduced orthoflavivirus database, DENV comprised 23.4% of sequences, substantially reducing the overrepresentation of this pathogen relative to the original database.

#### 3.1.3. Clustering and reduction of the *Alphavirus database*

At a major cluster ID of 0.70, species were not clearly delineated. At 0.75, most *Alphavirus* species formed distinct clusters, with the exception of *A. madariaga*, which clustered with the majority of EEEV sequences, and *A. tonate*, *A.* mucambo and a subset of VEEV sequences, which clustered together (**Table S8**). CHIKV formed two clusters; VEEV formed eight clusters, with most sequences contained in one cluster; EEEV formed four clusters, again dominated by a single major cluster; WEEV formed three clusters, with the majority of sequences contained within one cluster, and SINV formed two distinct clusters (**Table S9**).

At a major cluster ID of 0.80, further subdivision was observed. CHIKV split into nine clusters, although most sequences were contained within two of these. VEEV formed 15 clusters, EEEV formed seven, WEEV four and SINV eight, with most sequences for each virus distributed across one or two dominant clusters. *A. madariaga* continued to group with a subset of EEEV sequences, whereas *A.tonate* separated from *A. mucambo*, which remained grouped with a single VEEV sequence.

Based on the observed species-level delineation, a major cluster ID of 0.80 was selected for subsequent analysis, as this threshold achieved the highest species level separation. However, several species may have been over-partitioned at this setting. At subclustering identity thresholds of 0.97 and 0.98, the database was reduced to 1,013 sequences in both cases, whereas at threshold 0.99, the database was reduced to 1,071 sequences. The difference was primarily attributed to an increased number of CHIKV subclusters. Given these results, a major cluster identity of 0.80 and a subcluster identity of 0.98 was selected for downstream alignment (**Table S10**). In this reduced *Alphavirus* database, CHIKV comprised 10.1% of sequences, substantially reducing the overrepresentation of this pathogen.

#### 3.1.4. Initial evaluation of CD-HIT and VSEARCH

Following the initial primer design (see section 3.3) for *Orthoflavivirus* and *Alphavirus*, clustering performance was evaluated by calculating silhouette scores from Mash distance matrices for clusters generated using VSEARCH and CD-HIT. For *Orthoflavivirus*, CD-HIT produced markedly lower silhouette scores than VSEARCH (**Figure S1 A**, **Table S11**), indicating lower intra-cluster cohesion and inter-cluster separation. For *Alphavirus*, CD-HIT yielded negative silhouette scores across all evaluated ID thresholds, indicating poorly separated clusters. In contrast, VSEARCH produced higher silhouette scores, indication more coherent clusters. However, VSEARCH required substantial computation time for each clustering and subclustering step.

### 3.2. Evaluation of clustering algorithms

#### 3.2.1. Implementation and evaluation of Vclust

To address the limitations observed with CD-HIT and VSEARCH, we evaluated the recently developed Vclust framework, which enables clustering based on tANI using a reusable all-vs-all similarity matrix. The six different clustering methods implemented in Vclust were evaluated across sequence identity thresholds of 0.50-0.90 in increments of 0.05 and compared with clusters generated using CD-HIT and VSEARCH (**Figure S1 B** and **C**). VSEARCH and Vclust were evaluated across the full range of identity thresholds, whereas CD-HIT was restricted to thresholds ≥0.80 due to limitations of the algorithm. Clustering performance was evaluated using silhouette scores calculated from both Mash and tANI-derived distance matrices.

For *Orthoflavivirus*, VSEARCH and Vclust algorithms generally produced higher silhouette scores than CD-HIT across identity thresholds were all three approached could be compared (**Figure S1 B**). VSEARCH performed particularly well at identity thresholds of 0.75-0.80, where its performance was comparable to Vclust-single. Across the Vclust algorithms, high silhouette scores were obtained over a range of identity thresholds, although the threshold associated with maximum performance varied among algorithms. No single Vclust algorithm consistently outperformed the others across the evaluated range.

A similar pattern was observed for *Alphavirus*, where Vclust generally performed comparably to VSEARCH and substantially better then CD-HIT (**Figure S1 C**). Using tANI-derived distances, VSEARCH and several Vclust algorithms achieved silhouette scores above 0.8 across multiple identity thresholds. Overall, the results indicate that several Vclust algorithms can produce similarly coherent cluster structures, with clustering performance influenced more strongly by the selected identity threshold than by the specific Vclust algorithm.

The choice of distance metric also influenced the calculated silhouette scores, particularly for *Alphavirus*. While Mash- and tANI-derived distances generally showed similar trends, Mash-based silhouette scores displayed greater variation across identity thresholds and were generally lower than the corresponding tANI-based scores for *Alphavirus*. This difference was likely due to the broader variation in genome length within the Alphavirus dataset. In contrast, tANI-derived distances produced more consistent silhouette scores across clustering approaches and thresholds. As Vclust inherently computes an ANI matrix, tANI-derived distances were subsequently used for silhouette scoring of Vclust-generated clusters, ensuring consistency between clustering and evaluation. For VSEARCH and CD-HIT generated clusters, Mash distance matrices were retained for silhouette scoring due to their substantially lower computational cost.

#### 3.2.2. Computation comparison

Runtime and peak memory usage were compared for the complete clustering workflow using the *Orthoflavivirus* database (**Figure S2**). Overall, Vclust showed substantially lower wall-clock runtime than VSEARCH, reducing the total workflow time by nearly an order of magnitude. Differences among the Vclust algorithms were minor, with wall-clock runtimes ranging from approximately 2h 20min to 2h 30min, compared with 22h 20min for VSEARCH.

In contrast, peak memory usage was considerably higher for the Vclust-based workflows (ca. 95-100 GB), relative to VSEARCH (ca. 10 GB). This increased memory usage does not appear to originate from the Vclust algorithms themselves, but rather from how the workflow handles and manipulates the all-vs-all ANI matrix during data import, intermediate storage, and repeated subclustering steps. The matrix is retained in memory to allow rapid generation of multiple subcluster partitions, which increases efficiency for repeated analysis at the cost of higher memory demand. Despite this trade-off, the overall computational performance of Vclust remains favorable, particularly when clustering must be performed repeatedly across multiple identity thresholds.

#### 3.2.3. Species-level separability

To assess the congruence between genomic clustering and formal species taxonomy, silhouette scores were calculated using species assignments of ICTV-classified sequences. For *Orthoflavivirus*, the species-level silhouette score was 0.594 when calculated using tANI-derived distances, indicating moderate-to-strong separation among recognized species. The corresponding analysis based on Mash distances yielded a lower silhouette score (0.428), consistent with the higher sensitivity of Mash to genome length variability. In contrast, *Alphavirus* displayed substantially higher species-level separability, with silhouette scores of 0.926 and 0.894, using tANI-derived and Mash distances, respectively, reflecting a strong genomic distinction among species within this genus.

While genome-based distances capture much of the underlying taxonomic structure, they do not account for non-genomic criteria used by ICTV, such as antigenic, host, vector or ecological association. Nevertheless, these results demonstrate that sequence similarity alone explains a large fraction of species level differentiation in both genera, particularly for Alphavirus, where genomic divergence closely parallels established taxonomic boundaries.

### 3.3. Application to primer design

#### 3.3.1. *Orthoflavivirus* primer design and experimental evaluation

The CD-HIT reduced *Orthoflavivirus* database was aligned and processed in varVAMP with consensus thresholds and 0.75, 0.80 and 0.85 and maximum ambiguity settings (max_ambig) of 3, 4, 5, and 6. Across all parameter combinations, the qPCR mode failed to produce valid primer-probe sets, whereas the single mode successfully generated primer pairs (**Table S12**). In total, five primer sets were generated (**Table 1**), all targeting the NS5 region.

**Table 1:** Primers designed using the primer design workflow for orthoflaviviruses.

| Primer name | Sequence (5'-3') | Size<sup>α</sup> | Threshold<sup>β</sup> | Orthoflavivirus<sup>γ</sup> | JE-C<sup>δ</sup> |
| --- | --- | --- | --- | --- | --- |
| pan-flavi_01_F | TGY RTB TAC AAC ATG ATG GG | 277 | 0.80 | 14853/17644 | 2571/3249 |
| pan-flavi_01_R | TGT CCC AKC CDG CDG TGT C |  |  |  |  |
| pan-flavi_02_F | MGD GCY ATM TGG TWC ATG TGG | 211 | 0.80 | 14855/17644 | 2571/3249 |
| pan-flavi_02_R | TGT CCC AKC CDG CDG TGT C |  |  |  |  |
| pan-flavi_03_F | TGY RTB TAC AAC ATG ATG GG | 275 | 0.75 | 15710/17644 | 3110/3249 |
| pan-flavi_03_R | TCC CAK CCD GCD GTG TCA TC |  |  |  |  |
| pan-flavi_04_F | MGR GCY ATM TGG TAC ATG TGG | 209 | 0.75 | 15705/17644 | 3110/3249 |
| pan-flavi_04_R | TCC CAK CCD GCD GTG TCA TC |  |  |  |  |
| pan-flavi_05_F | GCY CTV AAC ACH YTC ACS AAC | 158 | 0.75 | 12872/17644 | 2720/3249 |
| pan-flavi_05_R | TCA TCY CCR CTS AYB RCC AT |  |  |  |  |

<sup>α</sup> Size is based on the PCR amplicon generated from the consensus sequence created in varVamp
<sup>β</sup>Consensus threshold for consensus sequence generation in varVamp
<sup>γ</sup>*In silico* analysis of the primer-set using a database with 17644 orthoflavivirus sequences
<sup>δ</sup>*In silico* analysis of the primer set using a database with 3249 sequences belonging to the JE serocomplex

*In silico* on-target prediction showed that primer sets pan-flavi 03 and pan-flavi 04 had the highest hit rates among all primer sets, both in regards to capturing *Orthoflavivirus* sequences, as well as members of the Japanese encephalitis (JE) complex. Primer set pan-flavi 05 displayed the lowest overall hit rate across *Orthoflavivirus* sequences, but showed greater coverage of the JE complex than pan-flavi 01 and pan-flavi 02. Based on these results, pan-flavi 03, pan-flavi 04, and pan-flavi 05 were selected for experimental evaluation. The selected primer sets were initially screened against *Orthoflavivirus* control samples (WNV-ctrl, USUV-ctrl, and JEV-ctrl). Pan-flavi 03 produced secondary amplification products in all reactions, including the no-template control, indicating nonspecific amplification (**Figure S3**). Pan-flavi 04 amplified all three targets, generating distinct melting-curve peaks consistent with specific amplification. In contrast, primer set pan-flavi 05 failed to amplify any of the three target controls.

Pan-flavi 04 was selected for further validation using a viral RNA panel comprising different *Orthoflaviviruses*, including WNV, USUV, JEV, TBEV and SLEV. The assay amplified majority of the samples (**Figure 2**); however, dual melting curve peaks were observed for USUV-Fr16 (**Figure 2 F**), WNV-Kunjin (**Figure 2 D**), WNV1-NY99 and WNV3 Rabensburg (**Figure 2 H**). In the reference assay, USUV-Fr16, WNV-Kunjin and WNV3 Rabensburg displayed Ct-values of 33.14, 30.57 and 27.21, respectively, whereas WNV1-NY99 had a substantially lower Ct-value at 20.61.

**Figure 2.**
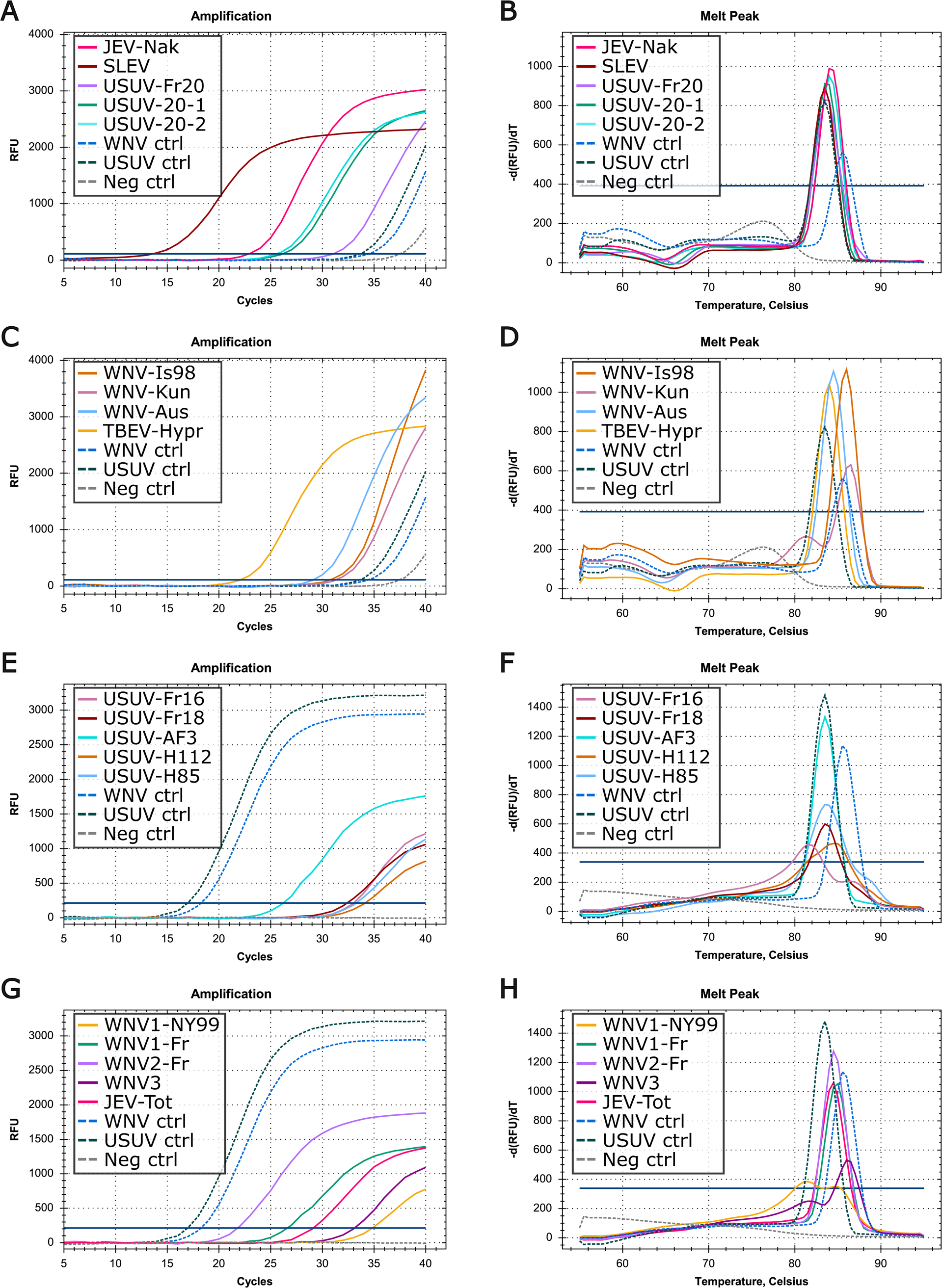
Validation of pan-flavi 04 on panel of orthoflaviviruses. A) Amplification curves for JEV Nakayama, SLEV, and USUV samples. B) Melting-curve profiles for JEV, SLEV and USUV samples. C) Amplification curves for TBEV and WNV samples. D) Melting-curve profiles for TBEV and WNV samples. E) Amplification curves for additional USUV samples. F) Melting-curve profiles for additional USUV samples. G) Amplification curves for JEV Tottori and additional WNV samples. H) Melting-curve profiles for JEV Tottori and additional WNV samples. WNV, USUV and negative controls as dashed lines. For A-D controls were diluted 10,000-fold. For E-G, controls were undiluted.

#### 3.3.2. *Alphavirus* primer design and experimental evaluation

The CD-HIT reduced *Alphavirus* database was aligned and processed using varVAMP with the same consensus thresholds and maximum ambiguity settings as described above. Similar to the *Orthoflavivirus* workflow, the qPCR mode failed to produce valid primer-probe sets across all parameter combinations, whereas the single mode successfully generated primer pairs (**Table S13**). In total, nine primer sets were generated (**Table 2**), targeting regions within nsP1, nsP2, nsP4 and E1.

**Table 2:** Primers designed using the primer design workflow for alphaviruses.

| Primer name | Sequence (5'-3') | Size<sup>α</sup> | Threshold<sup>β</sup> | On-target<sup>γ</sup> |
| --- | --- | --- | --- | --- |
| pan-alpha_01_F | KCN ATG ATG AAR TCH GGN ATG TT | 207 | 0.9 | 4194/4270 |
| pan-alpha_01_R | TYT TNA CYT CCA TRT TVA NCC A |  |  |  |
| pan-alpha_02_F | AAR TTY GGN GCB ATG ATG AA | 216 | 0.85 | 4260/4270 |
| pan-alpha_02_R | TYT TNA CYT CCA TRT TVA NCC A |  |  |  |
| pan-alpha_03_F | AAR TTY GGV GCB ATG ATG AA | 216 | 0.8 | 4260/4270 |
| pan-alpha_03_R | TYT TBA CYT CCA TRT TSA NCC A |  |  |  |
| pan-alpha_04_F | AAR TTY GGV GCB ATG ATG AA | 216 | 0.75 | 4260/4270 |
| pan-alpha_04_R | TCT TBA CYT CCA TGT TSA BCC A |  |  |  |
| pan-alpha_05_F | GAK TAC ATY ACH TGC RAV TAC AA | 156 | 0.75 | 3098/4270 |
| pan-alpha_05_R | ART RKG CBC CKC CCC ACA T |  |  |  |
| pan-alpha_06_F | CCD AAR CAR TGY GGM TTC TT | 267 | 0.75 | 3176/4270 |
| pan-alpha_06_R | TGY TTM ACC CAB CCK CKG AA |  |  |  |
| pan-alpha_07_F | TTY GAY ACH ACS CCR TTC AT | 267 | 0.75 | 3849/4270 |
| pan-alpha_07_R | AAY ACV GAH GGH AGR TGC CA |  |  |  |
| pan-alpha_08_F | AAT GAC CAT GCW AAY GCY AGA GC | 154 | 0.75 | 3017/4270 |
| pan-alpha_08_R | ATB GGR CAR AYR CAR TGG TA |  |  |  |
| pan-alpha_09_F | TRG AKT ACA TYA CHT GCR AVT ACA | 158 | 0.75 | 3168/4270 |
| pan-alpha_09_R | ART RKG CBC CKC CCC ACA T |  |  |  |

<sup>α</sup> Size is based on the PCR amplicon generated from the consensus sequence created in varVamp
<sup>β</sup>Consensus threshold for consensus sequence generation in varVamp
<sup>γ</sup>*In silico* analysis of the primer-set using a database with 4270 full-length alphavirus sequences

Primer sets pan-alpha 02, pan-alpha 03, and pan-alpha 04 targeted the same genomic region, and differed only in primer degeneracy. Pan-alpha 01 targeted a similar region, although the position of its forward primer differed from those of pan 02-04, whereas the reverse primers targeted similar positions. The forward primers of pan-alpha 05 and pan-alpha 09 also showed substantial positional overlap. *In silico* on-target prediction revealed considerable variation in predicted coverage among the designed primer pairs. Pan-alpha 01-04 and 07 exhibited the highest on-target hit rates, capturing 90.14%-99.77% of the *Alphavirus* database, whereas the remaining primer pairs exhibited lower hit rates ranging from 70.66%-74.38%. Based on the combined design characteristics and the *in-silico* predictions, pan-alpha 01, pan-alpha 02, and pan-alpha 05-09 were selected for experimental validation.

The selected primers pairs were initially screened against VEEV and CHIKV control samples by RT-qPCR. Pan-alpha 08 amplified both targets and produced distinct melting-curve peaks consistent with specific amplification, whereas the remaining primer pairs failed to produce the expected amplification products (**Figure S4**). Pan-alpha 08 was subsequently evaluated against a broader panel comprising VEEV, EEEV, and WEEV samples. The assay successfully amplified all samples in the panel, producing distinct melting-curve peaks consistent with specific amplification (**Figure 3**).

**Figure 3.**
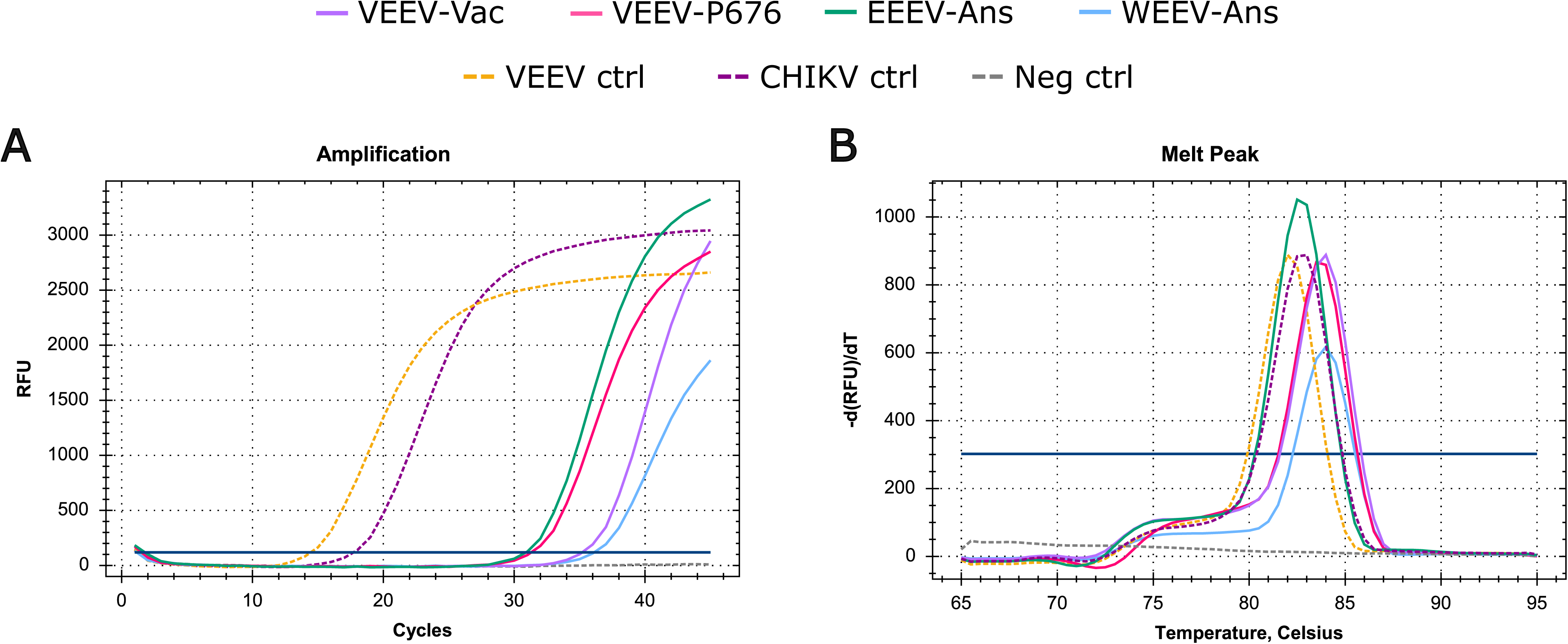
Validation of pan-alpha 08 on panel of alphaviruses. A) Amplification curves. B) Melting-peak profiles. VEEV, CHIKV and negative control as dashed lines. Samples were undiluted.

### 3.4. Reanalysis using Vclust-based clustering

#### 3.4.1. Analysis of Vclust single-linkage clustering

Given the lower clustering performance observed for CD-HIT in the silhouette score analyses, the clustering workflow was repeated using Vclust single-linkage clustering. Both databases were clustered using major cluster ID thresholds of 0.70, 0.75, 0.80, 0.85, and 0.90; subcluster ID thresholds of 0.96, 0.97, 0.98, and 0.99; and a target of *n*=100 sequences per major cluster.

For *Orthoflavivirus*, the highest silhouette score was obtained at major cluster ID 0.80 (score=0.807), indicating an optimal balance between intra-cluster cohesion and inter-cluster separation. Consistent with the silhouette scores, clustering at 0.80 achieved the most balanced partitioning, separating previously mixed taxa while maintaining species integrity and avoiding excessive fragmentation at higher thresholds. At lower thresholds, several species were not fully separated. At 0.70, TBEV, *O. langatense*, *O. loupingi*, and *O. omskense* clustered together; *O. bagazaense*, *O. israelense*, and *O. ntayaense* formed a joint cluster; *O. sepikense* and *O. wesselbronense* were grouped together; and *O. koutangoense* clustered with WNV. With increasing thresholds, these mixed clusters gradually resolved. At 0.75, *O. langatense* separated into its own cluster, whereas *O. omskense* remained grouped with TBEV and *O. loupingi* until 0.85. Even at 0.90, *O. bagazaense* and *O. israelense* continued to cluster together, and two TBEV sequences continued to group with the majority of *O. loupingi* sequences. *O. sepikense* and *O. wesselbronense* separated at 0.80, and at 0.85, *O. koutangoense* formed a distinct cluster together with a single residual WNV sequence. Cluster fragmentation increased within major species groups at higher thresholds. DENV formed three clusters at 0.70, six at 0.75 and 0.80, nine at 0.85 and twelve at 0.90, although four dominant clusters consistently contained the majority of the sequences. WNV expanded from two clusters at 0.75 to ten at 0.90, dominated by two principal groups. JEV and YFV each split into two clusters at 0.80 and 0.85, and four at 0.90, again with most sequences concentrated in one or two dominant clusters.

For this genus, overclustering penalties calculated for the optimal major cluster ID (0.80) were 0 for subcluster IDs 0.96 and 0.97, indicating that the number of subclusters remained within or below the target of 100 sequences. Slight overclustering was observed at 0.98 (weighted penalty = 0.007; mean penalty = 0.003), and more substantial overclustering at 0.99 (weighted penalty = 1.179; mean penalty = 0.061). The weighted score reflects the impact of overclustering in large clusters, whereas the mean score represents the proportion of clusters exceeding the target, providing an overall measure of its extent.

For *Alphavirus*, the highest silhouette score was obtained at major cluster ID 0.85 (score=0.844), although similar high scores (0.820-0.842) were observed at IDs of 0.70, 0.75, and 0.80, indicating well-separated clusters. At 0.70, several species were not fully separated: *A. eastern* and *A. madariaga* formed a joint cluster, as did *A. getah* with *A. rossriver*, and *A. highlandsj* with *A. western*. Increasing the ID threshold led to progressive resolution of these groups. At 0.75, *A. highlandsj* and *A. western*, as well as *A. getah* and *A. rossriver* separated into distinct clusters, while *A. sindbis* split into two major clusters. At 0.80, most *A. eastern* sequences formed an independent cluster, and *A. venezuelan* divided into two principal groups, and CHIKV split into a major and relatively minor cluster. At 0.85, clustering exhibited strong taxonomic coherence, with very little mixed-species clusters and minimal overfragmentation. Mostly isolated singleton sequences deviated from their primary groups (e.g. *A. venezuelan, A. sindbis*, and *A. eastern*), indicating minor sequence divergence within otherwise well-defined taxa. At 0.90, slight overfragmentation was observed, consistent with the lower silhouette score, with additional small clusters emerging within several (e.g. *A. venezuelan*, *A. sindbis*, CHIKV, and *A. eastern*). Together, these observations confirm that 0.85 represents the optimal major cluster threshold for Alphavirus, capturing high taxonomic coherence while minimizing both species mixing and overfragmentation. Overclustering penalties for all Alphavirus analyses were 0 across subcluster ID thresholds (0.96-0.99), indicating that the number of subclusters remained within the target and that no excessive fragmentation occurred.

#### 3.4.2. Primer redesign

Reduced databases were generated using the clustering parameters selected above. For *Orthoflavivirus*, the database generated using a major cluster ID of 0.80 and subcluster ID of 0.97 contained 1,864 sequences, of which approximately 22% belonged to DENV, compared with more than 70% in the original database) (**Table S14**). For *Alphavirus*, the database reduced with major cluster ID 0.85 and subcluster ID 0.99, contained 1225 sequences, with approximately 9.3% derived from CHIKV (reduced from 56%) (**Table S15**).

For *Orthoflavivivirus*, the qPCR mode failed to produce valid primer-probe sets at consensus threshold of 0.75, and only the single mode was subsequently applied. The single mode was successful at thresholds 0.75 and 0.80, but failed at 0.85, yielding three primer sets in total (**Table S16**). The forward primers pan-flavi-vclust_01_F and pan-flavi-vclust_02_F were identical to each other and matched the previously designed pan-flavi_01F and pan-flavi_03_F, respectively. The reverse primer pan-flavi-vclust_01_R differed from pan-flavi_01_R and pan-flavi_02_R at a single nucleotide, containing D (A/G/T) instead of K (G/T). Similarly, pan-flavi-vclust_02R differed from pan-flavi_03_R and pan-flavi_04_R at a single position, containing R (A/G) instead of D (A/G/T). Pan-flavi-vclust_03 differed from pan-flavi_05 at a single nucleotide position in both the forward and reverse primers, containing W (A/T) instead of Y (C/T) in the forward primer and G instead of R (A/G) in the reverse primer.

Consistent with the high sequence similarity between the original and redesigned primer sets, only minor differences were observed in the predicted on-target coverage. Compared with their closest CD-HIT-derived counterparts, the number of on-target hits increased from 14,853/17,644 to 14,859/17,644 for pan-flavi-vclust_01, from 14,853/17,644 to 15,659/17,644 for pan-flavi-vclust_02, and from 12,872/17,644 to 12,882/17,644 for pan-flavi-vclust_03.

For *Alphavirus*, the qPCR mode also failed to generate primers-probe sets at a consensus threshold of 0.75, and only the single mode was used thereafter. Primer design in the single mode was successful at thresholds 0.75 and 0.80, but failed at 0.85, producing four primer sets in total (**Table S16**). Pan-alpha-vclust_01 and pan-alpha-vclust_04 targeted a region within nsp4 that was not represented among the previously designed primer sets. These primer pairs were nearly identical, differing at a single nucleotide in the reverse primer, V (A/C/G) vs M (A/C). In contrast, the reverse primers pan-alpha-vclust_02_R and pan-alpha-vclust_03_R, targeting nsp2, were identical and differed from the previously designed pan-alpha_06_R by a single nucleotide, containing B (C/G/T) instead of K (G/T). Their forward primers overlapped but were shifted relative to one other.

Consistent with the limited overlap between the original and redesigned *Alphavirus* primer sets, substantial differences were observed in the primer prediction analysis. The redesigned primer sets exhibited on-target hit rates ranging from 77.94%-81.57% compared with 70.66%-99.77% for the original designs.

Overall, primer design based on the Vclust single reduced database for *Orthoflavivirus* showed high concordance with previously generated CD-HIT based designs, mostly differing only by a few degenerate positions. This agreement is consistent with the comparatively better clustering performance of CD-HIT for *Orthoflavivirus* relative to *Alphavirus*. In contrast, for *Alphavirus*, the concordance between Vclust single and CD-HIT based primer sets was low, with minimal sequence overlap and differing target regions, reflecting the greater impact of clustering structure on primer design outcomes for this genus.

#### 3.4.3. Experimental validation of redesigned primers

Given the high similarity between the *Orthoflavivirus* primer sets generated from CD-HIT and Vclust-reduced databases, no additional experimental evaluation of the redesigned primers was performed. In contrast, the greater differences observed between the original and redesigned *Alphavirus* primer sets prompted experimental validation of the Vclust-derived designs.

Accordingly, all four pan-alpha-vclust primer sets were initially screened against VEEV and CHIKV control samples by RT-qPCR. Pan-alpha-vclust 02, Pan-alpha-vclust 03, and Pan-alpha-vclust 04 amplified both controls, whereas pan-alpha-vclust 01 failed to amplify the CHIKV control (**Figure S5**). The three primer pairs that amplified both controls were subsequently evaluated against the same viral panel comprising VEEV, EEEV, and WEEV samples. Pan-alpha-vclust 04 amplified all samples in the panel and produced distinct melting curve peaks consistent with specific amplification (**Figure S6**). In contrast, pan-alpha-vclust 02 failed to amplify VEEV-Vac and VEEV-P676, whereas pan-alpha-vclust 03 failed to amplify VEEV-Vac.

## 4. Discussion

Public viral genome databases are often dominated by a small number of extensively sampled species, whereas other taxa remain comparatively underrepresented. The workflow presented here was developed to mitigate these sampling biases while retaining representative genetic diversity for downstream primer design. Application of the workflow to the genera *Orthoflavivirus* and *Alphavirus* demonstrated that substantial reductions in database size could be achieved while maintaining the genetic diversity necessary for robust primer development. Database sizes were reduced from 17,644 to 1,864 sequences for *Orthoflavivirus* and from 4,270 to 1,225 sequences for *Alphavirus*, while substantially reducing the dominance of highly represented species such as DENV and CHIKV, respectively.

The workflow was initially implemented using CD-HIT and VSEARCH, as these tools provide a straightforward framework for identity-based clustering and are widely used for large-scale sequence reduction (Ju et al., 2025). During the initial stages of workflow development, clustering outcomes was assessed qualitatively through manual inspection of cluster composition and species separation. This approach indicated that both clustering methods could generate reduced databases suitable for downstream primer design but provided no quantitative basis for comparing clustering quality or selecting clustering parameters. As the project progressed, however, it became apparent that a more objective framework was needed to compare clustering strategies and identify optimal clustering parameters. This led to the introduction of silhouette score analysis based on genome distance matrices to provide an objective measure of intra-cluster cohesion and inter-cluster separation and to facilitate systematic comparison of clustering strategies and identity thresholds.

Application of this framework revealed substantial differences in clustering performance that were not readily apparent from qualitative assessment alone. In particular, CD-HIT consistently produced lower silhouette scores than VSEARCH for both *Orthoflavivirus* and *Alphavirus*, with particularly poor performance for the latter. These results indicate that CD-HIT generated clusters that were less internally coherent and less distinct from one another, resulting in a less effective representation of the underlying genetic diversity. The particularly poor performance observed for *Alphavirus* is consistent with the broader range of genome lengths represented within this dataset, suggesting that sequence length heterogeneity may have contributed to the reduced clustering performance observed for CD-HIT.

Despite its superior clustering performance, the computational requirements of VSEARCH limited its practicality when multiple major clustering and subclustering thresholds were evaluated. As each clustering step required a separate sequence comparison, runtime increased substantially with the number of parameter combinations explored. This limitation motivated the evaluation of the recently developed Vclust framework, which enables clustering across multiple thresholds using reusable all-versus-all ANI matrix.

The silhouette score analysis demonstrated that Vclust consistently produced cluster structures comparable to those obtained with VSEARCH. Across the evaluated algorithms, clustering performance was influenced more strongly by the selected identity threshold than by the specific clustering method employed. Most Vclust algorithms converged on similar silhouette scores at intermediate ANI thresholds, indicating that the underlying biological structure of the datasets was largely captured robustly regardless of the clustering algorithm used. Although no single Vclust algorithm consistently outperformed the others, Vclust-single was selected for subsequent analyses because it provided a favorable balance between species-level separation and within-species fragmentation at intermediate ANI thresholds. The resulting clusters broadly corresponded to the expected species-level relationships while avoiding excessive fragmentation of closely related sequences.

An additional advantage of the Vclust framework was the availability of ANI-based distances for cluster evaluation. Mash distances provided a rapid approximation of genomic similarity and enabled practical evaluation of CD-HIT and VSEARCH clustering results, but showed greater sensitivity to variation in genome length and completeness. In contrast, tANI-derived distances produced more stable clustering patterns and provided a distance metric that was directly aligned with the clustering procedure itself. This reduced the disconnect between cluster generation and cluster evaluation, allowing more robust comparison of clustering thresholds.

The computational benchmarking highlighted an important trade-off between runtime and memory consumption. Compared to VSEARCH, Vclust reduced total workflow runtime by nearly an order of magnitude, making it substantially more practical for iterative exploration of clustering parameters and repeated workflow execution. This improvement was achieved at the cost of increased memory consumption associated with the storage and manipulation of the all-versus-all ANI matrix. Nevertheless, in workflows with recurring evaluation of multiple clustering thresholds, the reduction in runtime is likely to outweigh the additional memory requirements, particularly on high-performance computing systems.

A central objective of the workflow was to facilitate the generation of representative sequence datasets suitable for consensus-based primer design. *Orthoflavivirus* and *Alphavirus* genome datasets were selected as proof-of-concept datasets because they represent genetically diverse and diagnostic relevant viral genera that are subject to substantial sampling biases in publicly available sequence repositories. By reducing the influence of heavily sampled taxa, while preserving broader genetic diversity, the clustering workflow generated more balanced datasets for downstream alignment and primer design. These reduced datasets could subsequently be processed by varVAMP without extensive manual curation or sequence selection, thereby streamlining the development of consensus-based primer sets while maintaining broad representation of genetic diversity. An important consideration when applying clustering-based database reduction is the representation of rare or highly divergent lineages. The workflow addresses this challenge by allowing users to preferentially retain specified accessions during representative sequence selection and by providing control over the number of sequences retained in the reduced database. Consequently, lineages of particular biological, epidemiological, or diagnostic importance can be preserved even when represented by relatively few genomes. Nevertheless, the contribution of rare variants to downstream consensus generation remains dependent on the selected clustering and reduction parameters, and adequate representation of highly divergent lineages may require adjustment of the target database size.

The combination of clustering-based database reduction, systematic exploration of varVAMP design parameters, and integrated reporting provides a semi-automated and reproducible framework for pan-primer development. Conventional pan-primer design often relies on manual selection of representative sequences and subjective identification of conserved target regions. In contrast, the workflow presented here provides a systematic approach to sequence reduction, evaluates multiple combinations of consensus thresholds and ambiguity settings, and consolidates the resulting candidate primer designs into a unified report for downstream evaluation. By reducing the manual decisions and documenting the parameter combinations evaluated, the workflow improves reproducibility and facilitates re-evaluation and updating of primer sets as new viral genomes become available.

The primer target regions identified in the workflow corresponded to conserved genomic regions shared across genetically diverse viral species. For both *Orthoflavivirus* and *Alphavirus*, the workflow independently identified primer-binding sites within genomic regions that have previously been targeted by published broad-range assays (Chao et al., 2007; Eshoo et al., 2007; Giry et al., 2017; Grywna et al., 2010; Johnson et al., 2010; Kuno et al., 1998; Patel et al., 2013; Pfeffer et al., 1997; Sánchez-Seco et al., 2001; Scaramozzino et al., 2001). This concordance with previously published assays suggests that the workflow is capable of identifying biologically meaningful conserved regions while reducing the influence of uneven sequence representation.

Interestingly, the influence of clustering strategy on primer design outcomes differed markedly between the two proof-of-concept datasets. For *Orthoflavivirus*, redesign using the Vclust-derived database produced primer sets that were highly similar to those obtained from the CD-HIT-derived database, suggesting that the identified primer-binding regions were relatively robust to changes in clustering methodology. In contrast, substantially different primer sets were obtained for *Alphavirus*, indicating that primer design outcomes may be more sensitive to how sequence diversity is represented within the reduced dataset. This difference may partly reflect the greater overall diversity of the *Alphavirus* dataset, including its broader range of genome lengths than the *Orthoflavivirus* dataset. Consequently, representative sequence selection appears to have had a greater influence on the downstream primer design for *Alphavirus* than for *Orthoflavivirus*. These findings highlight that the consequences of clustering choices may not be uniform across viral groups and should therefore be evaluated in the context of the sequence diversity and composition of the dataset.

The results also highlight several challenges associated with consensus-based primer design. Although the workflow successfully generated candidate assays for both genera, the frequent inability of the qPCR mode to generate valid primer-probe assays demonstrates the difficulty of identifying sufficiently conserved target regions across genetically diverse viral groups. Whereas primer-only assays require two conserved primer-binding sites, primer-probe assays additionally require a suitable conserved probe-binding region. As sequence diversity increases, the likelihood of simultaneously identifying conserved primer and probe targets correspondingly decreases, potentially making genus-level assay design more challenging for probe-based than for primer-only formats. This observation is perhaps unsurprising given that varVAMP was originally developed for primer design within comparatively constrained target groups, whereas the present study applied the tool at the genus level across highly heterogeneous datasets. In this context, conserved target regions must span multiple species rather than a single species or lineage, imposing greater constraints on the sequence conservation available for primer and probe design.

Furthermore, broad inclusivity must be balanced against analytical performance. Primers designed to maximize coverage genetically diverse targets often require increased degeneracy, which may reduce amplification efficiency, increase the likelihood of non-specific amplification, and complicate assay optimization. Therefore, experimental validation remains essential, as predicted *in silico* coverage does not necessarily translate into optimal analytical sensitivity and specificity under laboratory conditions.

Given the extensive sequence diversity present within both *Orthoflavivirus* and *Alphavirus*, future optimization efforts could explore primer mixtures composed of multiple less-degenerate primer sets rather than relying on a single highly degenerate assay. While varVAMP aims to maximize inclusivity through the design of a single degenerate primer set, an alternative strategy would be to replace highly degenerate positions with mixtures of individual primers targeting the same genomic region. Such primer cocktails could retain broad sequence coverage while reducing the level of degeneracy within individual primers, which could potentially improve amplification efficiency, analytical sensitivity, and overall assay robustness. Future studies should therefore evaluate the relative performance of highly degenerate primers and primer mixtures to determine the optimal balance between broad inclusivity and analytical performance for broad-range pathogen detection.

Although demonstrated here using *Orthoflavivirus* and *Alphavirus*, the workflow is not inherently specific to either genus and is designed to be applicable to other genetically diverse viral groups for which representative sequence selection and consensus-based primer design are required. By combining clustering-based sequence reduction with systematic exploration of primer design parameters, the workflow provides a scalable and reproducible framework for broad-range primer design that reduces reliance on manual sequence selection while retaining representation of genetic diversity.

The resulting representative datasets facilitate the identification of conserved primer-binding regions suitable for assay development and can be updated as additional genomic data become available. By supporting the development of broadly reactive assays from representative sequence collections, the workflow provided a framework for design of pathogen-agnostic detection strategies, enhances the capacity for broad-range pathogen detection and improves preparedness for emerging, divergent, or previously unrecognized viruses. Its semi-automated design of the workflow also facilitates systematic re-evaluation of candidate broad-range assays as sequence databases expands, providing a flexible approach that may support surveillance, outbreak response, and future One Health preparedness efforts.

## Supplementary

**Table S1:** *Orthoflavivirus* samples

**Table S2:** *Alphavirus* samples

**Table S3:** *Orthoflavivirus* database

**Table S4:** *Alphavirus* database

**Table S5:** *Orthoflavivirus* shared clusters

**Table S6:** *Orthoflavivirus* split species clusters

**Table S7:** *Orthoflavivirus* reduced database

**Table S8:** *Alphavivirus* shared clusters

**Table S9:** *Alphavirus* split species clusters

**Table S10:** *Alphavirus* reduced database

**Table S11:** Silhouette scores

**Table S12:** *Orthoflavivirus* varVamp run status

**Table S13:** *Alphavirus* varVamp run status

**Table S14:** *Orthoflavivirus* vclust reduced database

**Table S15:** *Alphavirus* vclust reduced database **Table S16:** Primer redesign

**Figure S1:** Comparison of clustering methods. A) Comparison of CD-HIT and VSEARCH for *Orthoflavivirus* and *Alphavirus* for major ids utilized in the initial clustering workflow using Mash distance matrix. B) Comparison of clustering methods for *Orthoflavivirus* using both Mash (solid) and tANI (dashed) distance matrices. C) Comparison of clustering methods for *Alphavirus* using both Mash (solid) and tANI (dashed) distance matrices.

**Figure S2:** Computational resource requirements. A) Computational benchmarking of clustering algorithms, CD-HIT and VSEARCH, applied to the *Orthoflavivirus* database across multiple identity thresholds. Each clustering run was performed in triplicate. Bars represent total runtime (yellow) and peak memory usage (blue). B) Computational benchmarking of complete clustering workflows comparing the six Vclust algorithms with VSEARCH. Each workflow was executed in triplicate, and mean values were used for comparison. Major clustering thresholds ranged from 0.70 to 0.90 (in 0.05 increments), and subclustering thresholds from 0.96 to 0.99 (in 0.01 increments). Bars show the total runtime (yellow) or peak memory (blue) usage for each workflow, subdivided into the distance matrix calculation (hatched) and the remaining clustering workflow (solid). All benchmarks were performed using 10 CPUs and 128 GB of RAM under identical SLURM configurations.

**Figure S3.** Screening of designed *Orthoflavivirus* primer sets. A) Amplification curves for primer set pan-flavi 03. B) Melting-curve profiles for pan-flavi 03. C) Amplification curves for pan-flavi 04. D) Melting-curve profiles for pan-flavi 04. E) Amplification curves for pan-flavi 05. F) Melting-curve profiles for pan-flavi 05.

**Figure S4.** Screening of designed *Alphavirus* primer sets. A) Amplification curves for primer set pan-alpha 01. B) Melting-curve profiles for pan-alpha 01. C) Amplification curves for pan-alpha 02. D) Melting-curve profiles for pan-alpha 02. E) Amplification curves for pan-alpha 05. F) Melting-curve profiles for pan-alpha 05. G) Amplification curves for pan-alpha 06. H) Melting-curve profiles for pan-alpha 06. I) Amplification curves for pan-alpha 07. J) Melting-curve profiles for pan-alpha 07. K) Amplification curves for pan-alpha 08. L) Melting- curve profiles for pan-alpha 08. M) Amplification curves for pan-alpha 09. N) Melting-curve profiles for pan- alpha 09.

**Figure S5.** Screening of redesigned *Alphavirus* primer sets. A) Amplification curves for primer set pan-alpha- vclust 01. B) Melting-curve profiles for pan-alpha-vclust 01. C) Amplification curves for pan-alpha-vclust 02. D) Melting-curve profiles for pan-alpha-vclust 02. E) Amplification curves for pan-alpha-vclust 03. F) Melting- curve profiles for pan-alpha-vclust 03. G) Amplification curves for pan-alpha-vclust 04. H) Melting-curve profiles for pan-alpha-vclust 04.

**Figure S6.** Validation of selected assays on panel of alphaviruses. A) Amplification curves for pan-alpha-vclust 02. B) Melting-curve profiles for pan-alpha-vclust 02. C) Amplification curves for pan-alpha-vclust 03. D) Melting-curve profiles for pan-alpha-vclust 03. E) Amplification curves for pan-alpha-vclust 03. F) Melting- curve profiles for pan-alpha-vclust 04. Samples were diluted 100-fold.

## Declarations Ethical approval

Not applicable.

## Consent for publication

Not applicable.

## Availability of data and materials

The datasets used and/or analyzed during the current study are available from the corresponding author on reasonable request.

The semi-automatic pan-primer design workflow is openly available on Github at https://github.com/Mailinnia/Semiautomatic_pan-primer_design. The version described in this study is permanently archived on Zenodo (Johnston, 2026)

## Authors’ contributions

**CMJ:** Conceptualization, Data curation, Formal analysis, Investigation, Methodology, Project administration, Software, Visualization, Writing – original draft, Writing – review and editing. **AM:** Methodology, Software, Writing – review and editing. **AMH:** Conceptualization, Supervision, Writing – review and editing. **LL:** Funding acquisition, Supervision, Writing – review and editing. **TBR:** Conceptualization, Funding acquisition, Project administration, Supervision, Writing – review and editing.

## Acknowledgements

We thank Jani Christiansen for her invaluable technical assistance throughout this study. We also thank Vithiagaran Gunalan and Thor Bech Johannesen for their valuable input during the initial discussions that helped shape the project approach.

