## Supplementary figures and images for "Efficient primer design for broad-range virus detection: A semi-automated workflow using sequence clustering and varVAMP"

### Supplementary figure S01

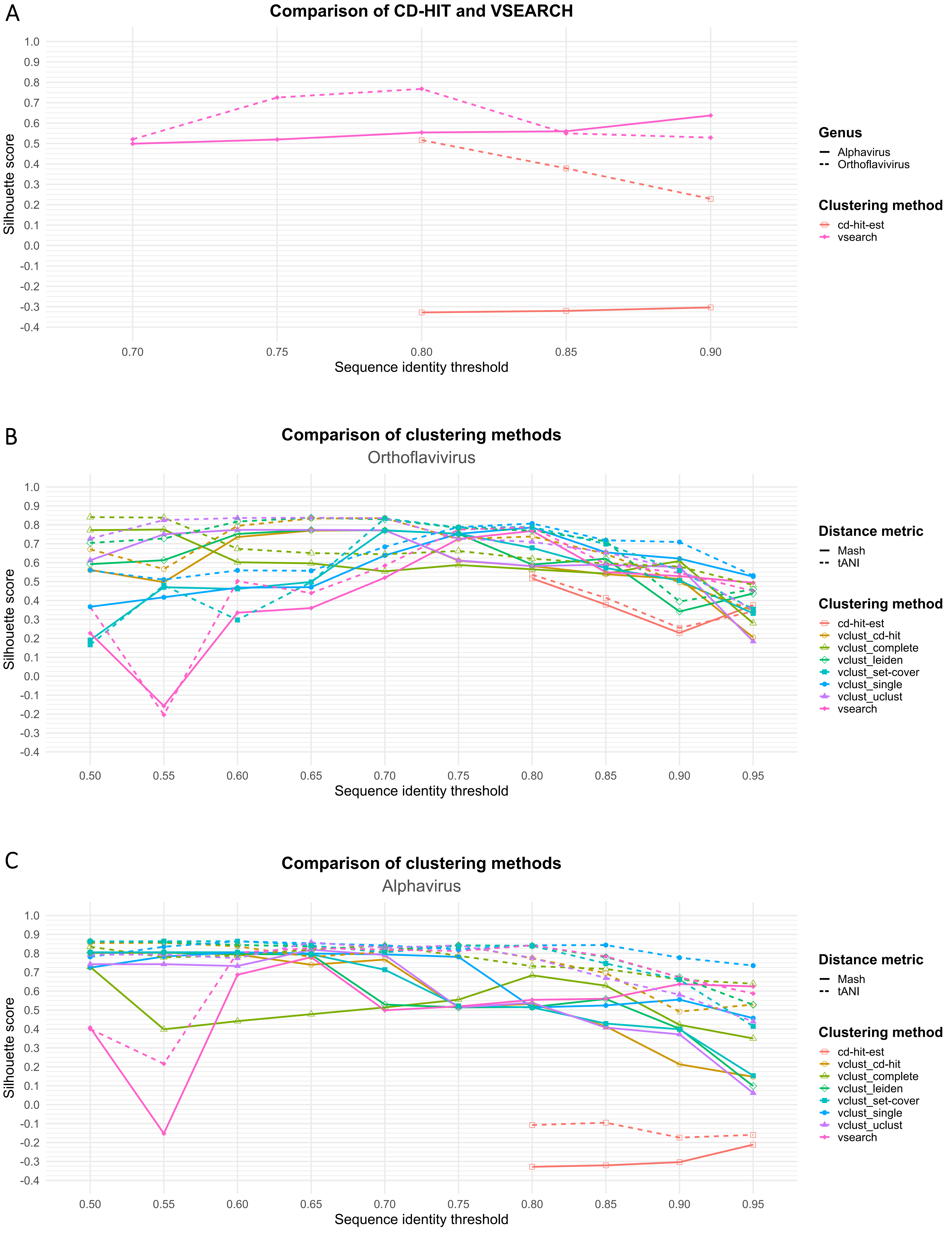

### Supplementary figure S02

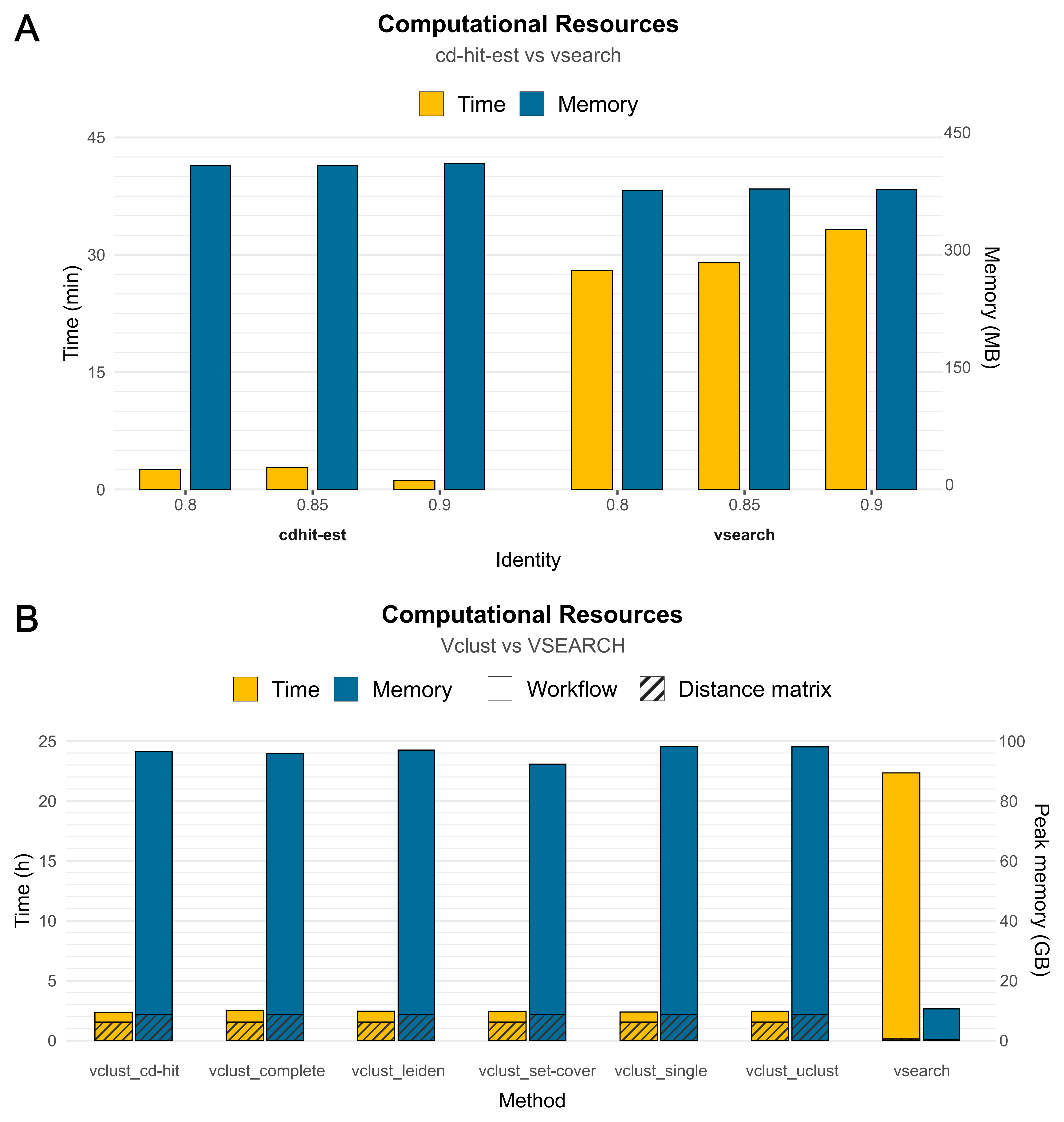

### Supplementary figure S03

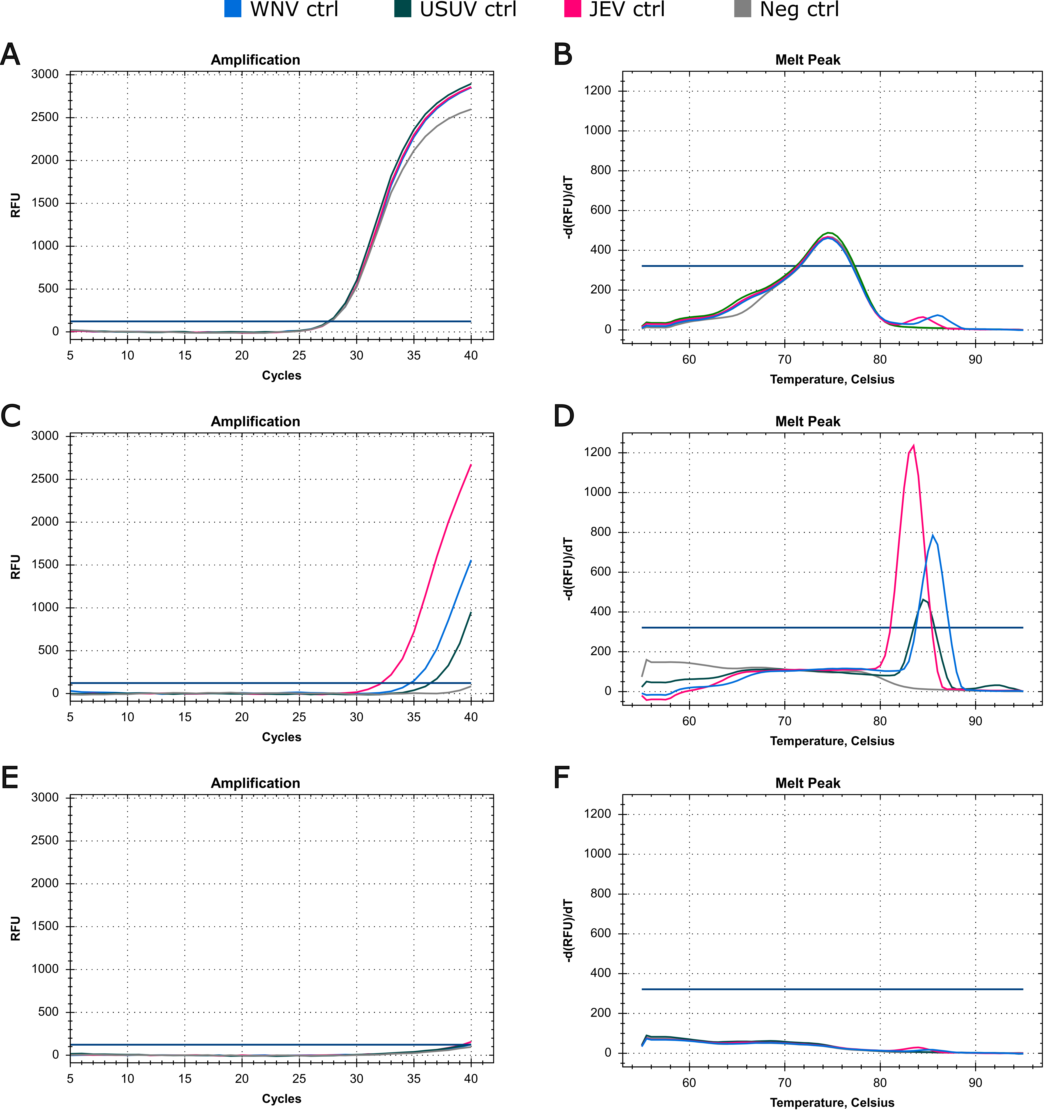

### Supplementary figure S04

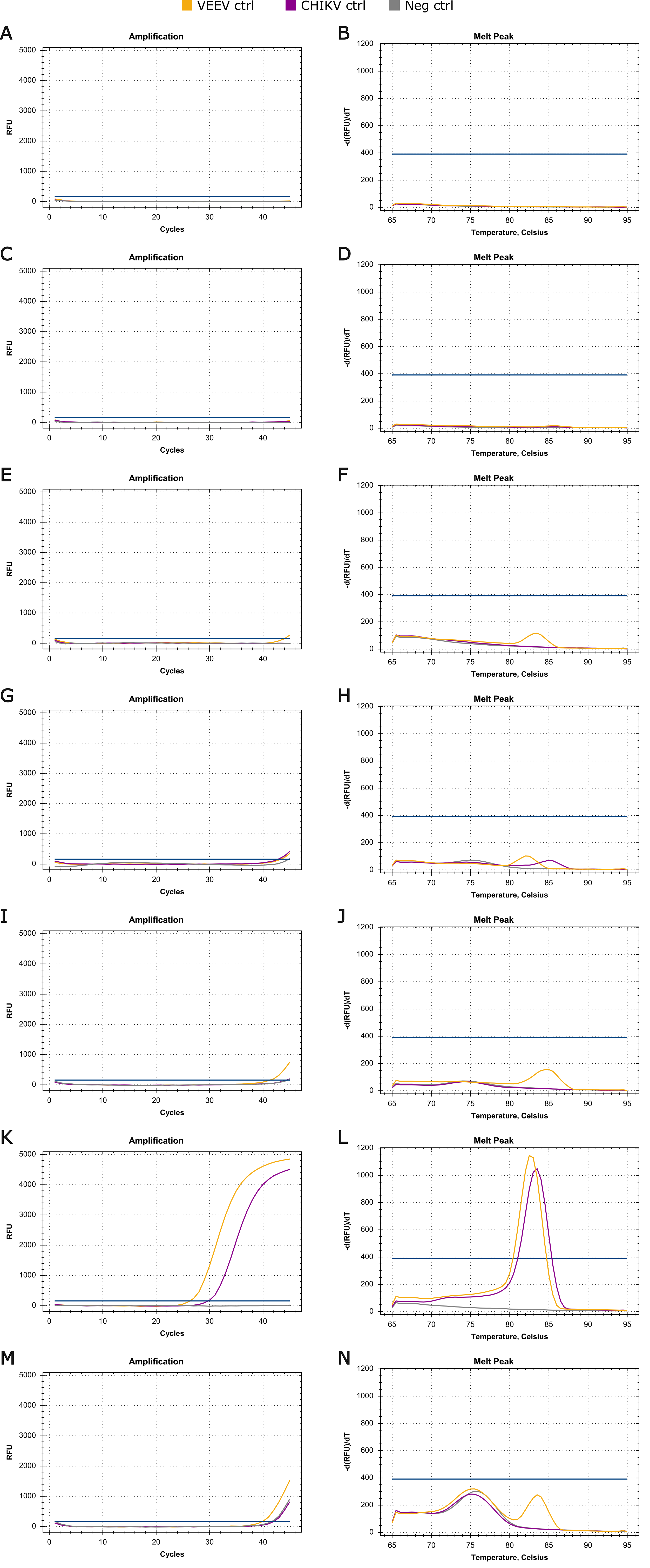

### Supplementary figure S05

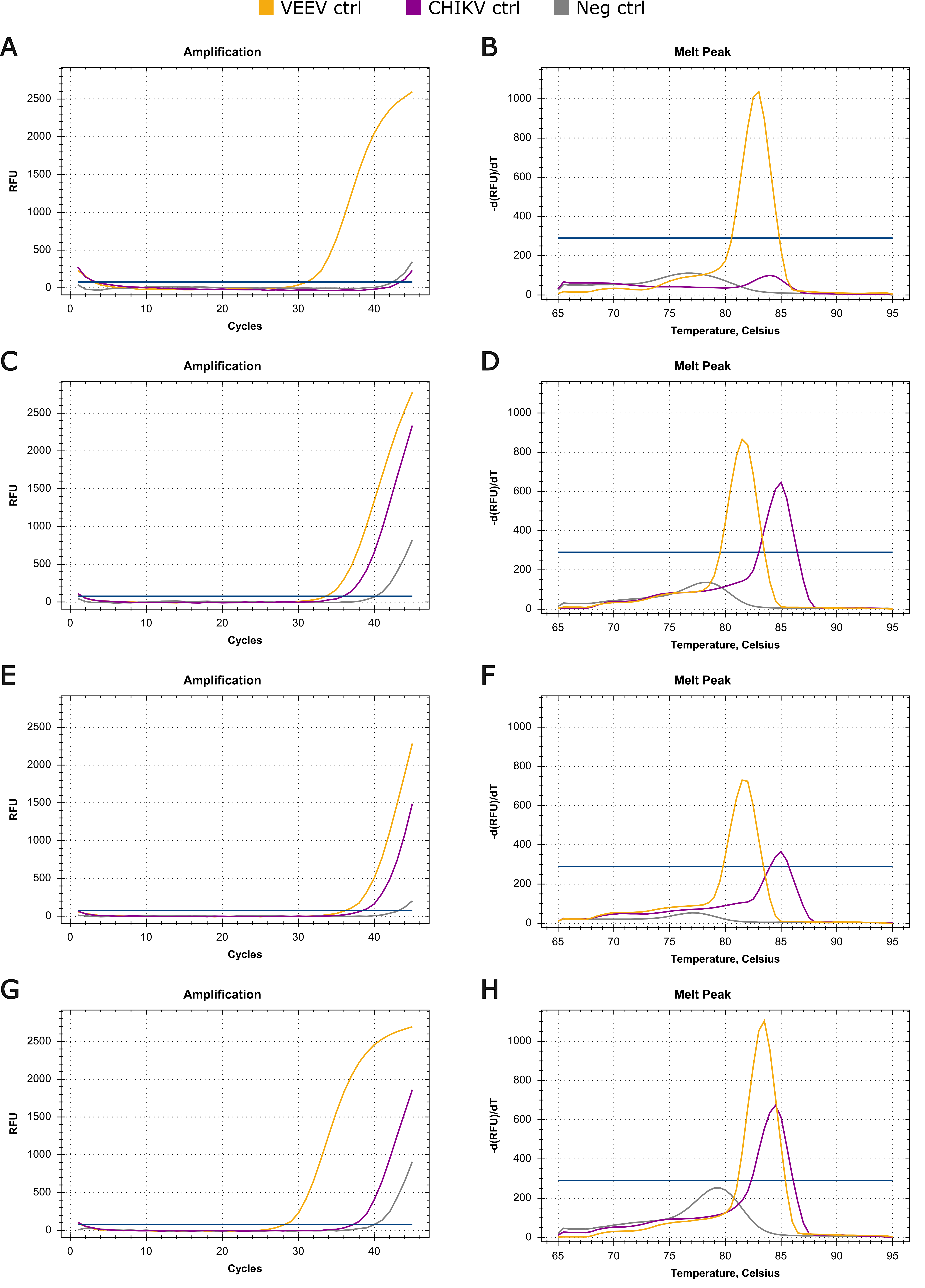

### Supplementary figure S06

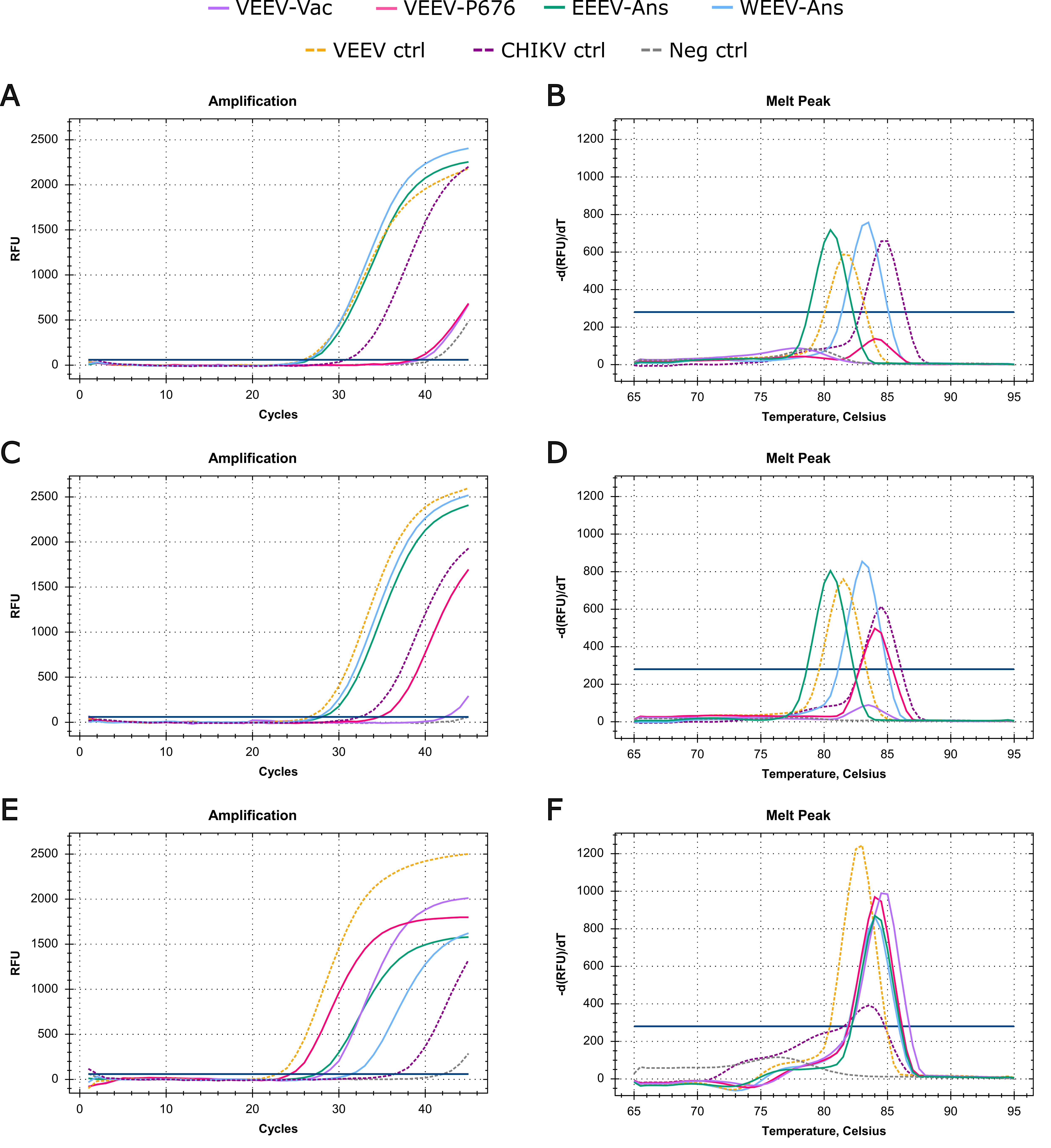
